# Integrating Data Across Oscillatory Power Bands Predicts the Seizure Onset Zone in Focal Epilepsies

**DOI:** 10.1101/2024.05.31.596825

**Authors:** Sean O’Leary, Anne-Cecile Lesage, Liliana Camarillo-Rodriguez, Oliver Zhou, Diosely C. Silveira, Jiefei Wang, Sameer A. Sheth, Joshua M Diamond, Michael S. Beauchamp, Zhengjia Wang, John F. Magnotti, Patrick J. Karas

## Abstract

Accurate identification of the seizure onset zone (SOZ) using intracranial electroencephalography (iEEG) remains challenging. Although diverse methods have leveraged spectral features to classify patient outcomes, few approaches focus on identifying individual electrodes within the SOZ or integrate a broad spectrum of frequency ranges within a single model. We developed an interpretable machine learning model that integrates power across delta, theta, alpha, beta, gamma, and high-gamma frequencies over time to identify the SOZ. For 1,511 electrodes implanted across 21 patients, we computed the mean spectral power in each frequency band for the first 20 seconds after seizure onset and analyzed the differences in power between SOZ and non-SOZ electrodes. In patients who were seizure-free after surgery (n = 14), electrodes within the SOZ showed significantly higher area under the curve (AUC) for mean power over time in the first 20 seconds after seizure onset compared to electrodes outside the SOZ in the alpha (p = 0.0272), beta (p = 0.0263), gamma (p = 0.0013), and high gamma (p = 0.0086) ranges. Additionally, electrodes within the SOZ in patients that became seizure-free after surgery had significantly higher AUC compared to equivalent electrodes in patients who did not become seizure-free after surgery (n = 7) in the gamma (p = 0.0145) and high gamma (p = 0.0024) power ranges. We trained a stacked random forest ensemble model using these features over time to label electrodes within the SOZ. Leave-one-out patient cross validation of the machine learning model yielded a 96.6% positive predictive value and 99.9% specificity for identifying electrodes within the SOZ. Our dataset included a diverse array of seizure onset patterns, which were all classified accurately by the model. A second model was trained to predict post-operative seizure freedom, yielding 95.2% accuracy for predicting seizure outcome based on a planned resection. This two-model design mirrors clinical workflow, first localizing SOZ electrodes to support surgical planning, then predicting outcome based on a surgical plan. An advantage of our interpretable machine learning approach is the ability to interrogate our models to understand how predictions are made. For electrode classification, the model weighed beta (0.66 ± 0.07), high gamma (0.54 ± 0.06), and delta (0.51 ± 0.06) power bands most heavily. Viewing the model’s frequency band weights over time reveals that the model identified a pattern resembling the “fingerprint of the epileptogenic zone”, reinforcing the importance of this dominant fundamental neurophysiologic signature of seizure onset.

## Introduction

Drug-resistant epilepsy affects approximately one third of people with epilepsy and is associated with significantly increased lifetime morbidity and mortality.^1–3^ Surgery is the most effective treatment for many of these patients. Effective procedures involve identifying and removing the site of seizure onset.^4^ Correctly defining this site of seizure onset, which we will refer to as the seizure onset zone (SOZ),^5, 6^ is of paramount importance to surgical success. When non-invasive diagnostic exams do not conclusively identify the SOZ, intracranial electrodes (iEEG) are implanted and the SOZ is identified by an epileptologist interpreting anatomo-clinico-electrical correlations of iEEG recordings.^7^ Epileptiform discharges are visually identified from iEEG recordings and correlated with their anatomic location and the patient’s clinical symptoms. Surgical outcomes (seizure freedom versus continuation of seizures) indicate whether the SOZ was correctly identified and removed, serving as a post-hoc confirmation. Using surgical outcomes as validation, various computational SOZ identification algorithms have been developed. These computational algorithms seek to identify and distill a subset of signal features from iEEG recordings to identify electrical SOZ biomarkers.

Many computational SOZ identification algorithms analyze iEEG power spectra at seizure onset, often focusing on singular frequency ranges such as high-frequency activity in beta^8, 9^ or gamma^10, 11^ bands, or slow rhythms in the delta, theta, or alpha ranges.^12, 13^ The epileptogenic index, for example, assesses the timing of changes in the ratio of energy between fast (beta and gamma) and slow (alpha and theta) power spectra.^14^ Another technique, the fingerprint, characterizes preictal spiking, early narrow-band fast activity and broad low-frequency suppression to define characteristic spectral patterns of the SOZ.^15^ A third method commonly referred to as DC-shift detects the co-occurrence of ictal direct current shifts (infraslow power below the delta band) and high-frequency oscillations to mark the SOZ.^16^ These diverse approaches highlight a complex interplay of neural activity across many frequency bands during the early ictal period. But changes in spectral power are dynamic in time, fluctuating in importance between different frequency ranges at different times.^13^ Moreover there is significant heterogeneity in broadly accepted seizure onset patterns between patients.^13, 17^ This variability necessitates adjusting computational methodologies to account for differences in seizure type and onset pattern.^18^ g

We propose an interpretable, supervised machine learning approach to develop a unified model considering iEEG seizure onset patterns across a broad frequency range (0.5-150 Hz). This model, named frequency range <u>e</u>xplorer of the <u>epileptogenic zone</u> (FREEZ), is more robust compared to existing methods that focus on a subset of spectral bands. The approach has two components, one model trained to label intracranial electrodes as within or not-within the SOZ, and a second model to predict patient outcomes given a proposed surgical resection. FREEZ consists of multiple random forest models that correlate the SOZ with spatiotemporal spectral power in each frequency band (delta, theta, alpha, beta, gamma, high-gamma). The individual random forest models are then integrated into an ensemble learning model.^19^ The resulting interpretable machine learning algorithm weighs spectral changes across time around the SOZ, deriving a dominant fundamental electrophysiological marker of the SOZ across power bands.

A critical hurdle facing adoption of computational SOZ localization algorithms is lack of validation, largely stemming from the difficulty of carefully critiquing and reproducing complex computational methods. To address this hurdle and make our algorithm easily available, we also created an opensource software package titled FREEZ^20^ available for free download along with web application demo: https://zenodo.org/records/11440403. The web application demo recreates Figure 1 so users can interact with data in the figure through a web GUI.

**Figure 1:**
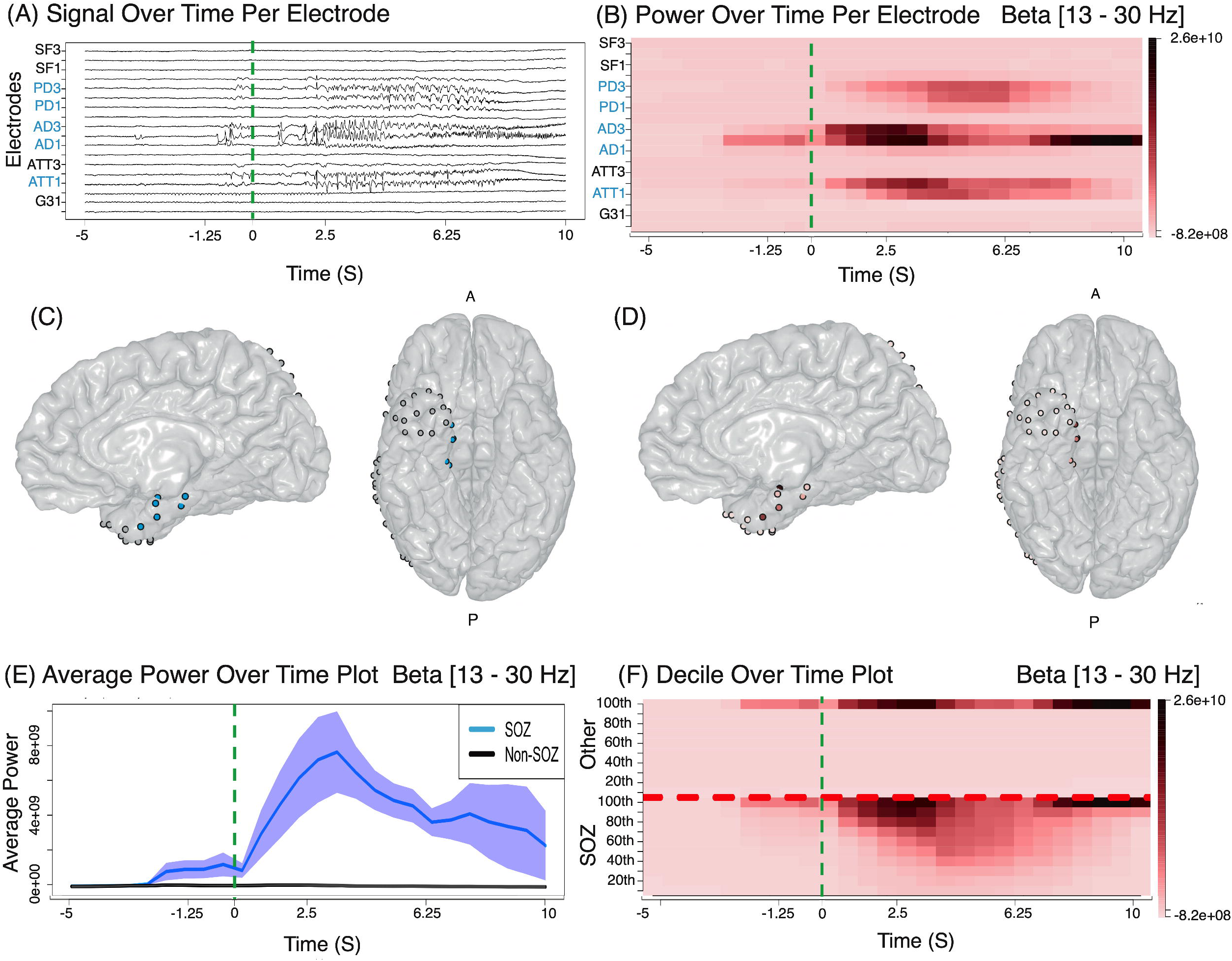
FREEZ module results on PT01. Plots are shown generated from the module displaying: (1A) Ictal intracranial EEG signal. (1B) Multitaper power result over time per electrode. (1C) 3D brain projection with marked SOZ channels. (1D) Projected multitaper power result over time in the beta frequency range (13–30 Hz). (1E) Average power over time line plot. (1F) Time plot of power percentiles from the 10th to the 100th (in steps of 10) for SOZ and non-SOZ channels. Data spans from -5 to 10 seconds around seizure onset, indicated by a green dotted line. Interactive FREEZ web application to view results displayed in this figure can be found here: https://rave-freez.azurewebsites.net/launcher/

## Material and Methods

### Patient Data and Selection

The Fragility multi-center dataset^8, 21^, available in part publicly on OpenNeuro, encompasses data from 35 patients distributed across four surgical epilepsy centers. The iEEG data is in the iEEG-BIDS format^22^ with signal data is in BrainVision Core Data Format (Brain Products GmbH, Gilching, Germany). Clinicians at each clinical facility labeled electrodes within the hypothesized SOZ. Electrode names and electrode classification were available, but electrode localization was not provided for all patients. The dataset features an event file with markers annotated by clinicians to indicate the timing of seizure onset/offset clinically and electrographically. Electrographic seizure onset times were used in this study and independently validated by D.C.S., a board-certified epileptologist with significant experience in surgical epilepsy. These seizures are marked according to standard procedures within the clinical epilepsy workflow. Additional details about the dataset are available.^8^

The following inclusion criteria were used: (1) patient underwent resection or laser ablation surgery guided by iEEG; (2) had at least one seizure recorded during intracranial monitoring, allowing for confirmatory testing of selected channels; (3) their recordings were at frequencies of 500 Hz or higher, suitable for spectral frequency analysis up to 150 Hz; and (4) detailed surgical outcomes, including Engel scores, were available. As in the study by Li *et al.* in which this data was originally published ^8^, we assume for the validation of our SOZ localization tools that electrodes annotated SOZ are inside the resected area for all patients and that they are a good estimate of the SOZ in case of seizure freedom outcome. When referring to the preoperative SOZ, we are referring to the annotated SOZ. Post-operative seizure freedom is used as an indicator of if the “true SOZ” was resected/ablated.

Seizure onset patterns for each patient were systematically analyzed according to the classification criteria established by Lagarde et *al.* ^28^, aiming to assess the accuracy of SOZ identification across a diverse array of seizure onset patterns. The protocol adhered strictly to the methodology outlined by Lagarde et *al.* ^28^. D.C.S categorized the onset pattern of all seizures. Visual analysis was conducted without the use of software filters, while spectral analysis utilized both the RAVE platform and the FREEZ module.^28^ The eight classified seizure onset patterns included A) low voltage fast activity (LVFA); B) preictal spiking with LVFA; C) polyspike bursts with LVFA; D) slow waves followed by LVFA; E) rhythmic slow spikes; F) sharp theta/alpha activity; G) beta sharp activity; and H) delta-brush.^28^

### iEEG data preprocessing in RAVE

Raw signal was preprocessed in RAVE 2.0.^28^ Data was epoched for seizure onset time and baseline signal. Electrodes with artifacts were removed from the analysis using visual inspection of the traces. Data were further notch filtered and then referenced using a common average reference (CAR) montage. These steps were performed using the RAVE 2.0 graphical user interface. Additional details about iEEG data preprocessing in RAVE 2.0 can be found in the RAVE 2.0 tutorial (https://rave.wiki/).

### Spectral spatiotemporal power analysis

For each electrode, we computed the mean spectral power for six frequency bands from the raw signal: delta (0.5-4 Hz), theta (4-8 Hz), alpha (8-13 Hz), beta (13-30 Hz), gamma (30-90 Hz), and high gamma (90–150 Hz) (**Fig. 1A** and **1B**). Calculations were performed using the multi-taper Fourier transform ^29^ method and computed using the multi-taper R library ^30^ (step size 0.5s, window length 2.5s). Power was estimated as power spectral density (PSD), with units of microvolts squared per hertz (μV²/Hz).^31^ These computations were performed across two epochs, the ictal interval (first 20 seconds after seizure onset) and the baseline period (10 second time interval starting 30 before seizure onset). Additional details about the multi-taper method and specific parameters are available in the *Supplementary Methods*. Clinical labels were then displayed overlayed on the reconstructed electrode implantation (**Fig. 1C**) and compared with the results of the mean spectral power (**Fig. 1D**).

### Statistical analysis of correlation between spectral power and SOZ

To analyze power change at seizure onset, mean spectral power output from multi-taper transform was exported for each electrode in .csv format for each of the six frequency ranges. The resulting time-frequency matrices were grouped based on electrode labels as SOZ *vs.* non-SOZ. For each timepoint within the analysis window, average power and standard error were computed for grouped SOZ electrodes and grouped non-SOZ electrodes at every frequency range. Electrode labels (SOZ *vs*. non-SOZ) were correlated to surgical outcome (seizure free vs. not seizure free) by calculating area under the curve (AUC) for these averaged power timeseries at every frequency range (**Fig. 1E**). To evaluate significant differences in AUC for each category, pairwise two-sample t-tests were conducted. A Bonferroni correction was applied to all p-values to adjust for multiple comparisons (24 pair-wise AUC comparisons).

Since each patient implantation requires a different number of electrodes, we computed the distribution of power across frequency bands in SOZ *vs.* non SOZ electrodes (using clinician labels) to predict post-operative seizure freedom outcomes.^8^ For each frequency band, we ranked the electrode power into deciles from 10-100 over the analysis time window for SOZ and non-SOZ electrode populations separately. Decile plots were created from this analysis (**Fig. 1F**).

### Ensemble machine learning method for multi-frequency analysis

Resulting power across all frequency bands was used to train two separate ensemble machine learning algorithms. One algorithm was trained to identify electrodes as within or not within the SOZ (*electrode classification*). A second algorithm was trained to predict post-operative seizure freedom based on removal of preselected electrodes (*patient outcome prediction*).

#### ELECTRODE CLASSIFICATION (SRFE-*electrode*)

To determine the spectral features that could distinguish between electrodes labeled as SOZ *vs.* not SOZ we used a stacked random forest ensemble (SRFE). This method builds a model to identify various interactions between spectral patterns present in the training data that correlate with the SOZ.

To create SRFE-*electrode*, we first trained six unique random forest models, one for each of the six frequency bands. Each model was trained with 100 decision trees using the seizure-free patient cohort, with clinician-determined SOZ electrode labeling as ground truth. We used the R H2O package ^32^ to compute the optimal coefficients (second-learning) needed to combine the six random forest models into the SRFE. The H2O R Library was chosen for ensemble learning training as it allows for packaged implementation, testing, and scaling of machine learning pipelines with a wide variety of models integrated together. The hyperparameters for SRFE-*electrode* included the selection of a Generalized Linear Model (GLM) as the metalearner algorithm.^33, 34^ Alpha is the regularization distribution parameter, we used the default parameter alpha equal to 0 (Ridge regression) to prevent overfitting and enhance generalization.^33, 34^ Lambda is the regularization strength parameter, we enabled lambda search to find the optimal parameter.^33, 34^ The SRFE model was then tested using leave-one-out cross validation. To convert prediction probabilities to binary SOZ labels (SOZ *vs.* non-SOZ) we tuned the model to prioritize specificity over sensitivity to avoid false positive SOZ labeling (reduce the potential for unnecessary tissue resection). This tuning was done by thresholding the model prediction probabilities to maximize true positives while maintaining positive predictive value (PPV) ≥ 0.95. Majority voting (≥ 50%) across different seizures for each patient was then used to select final electrode SOZ labeling.^15^

We additionally compared the SRFE method with two other machine learning methods using variations of random forest models. In one model, we trained a “simple random forest” on features consisting of all unique analysis time windows for all six frequency ranges. In the second case, we trained a unique random forest on each frequency range, as performed in the SRFE, but combined them by averaging their prediction results (probability that an electrode is in the SOZ) without using a SRFE model “averaged random forest”. Both random forest models were tuned similarly to the tuning of the SRFE model described above. Each model’s performance (simple random forest, averaged random forest, and SRFE) was evaluated with the following metrics: accuracy, specificity, sensitivity, PPV, and negative predictive value (NPV) to compare the predicted outcome with the ground truth electrode labels. For testing of all models, leave-one-out patient cross validation was employed. For patients with multiple seizures, we applied majority voting (≥ 50%) across ictal events to determine the final prediction.

#### PATIENT OUTCOME PREDICTION (SRFE-*outcome*)

Using the deciles calculated across frequency bands for SOZ vs. non-SOZ electrodes (**Fig. 1F**), we first normalized these decile values in each time window between 1 and 10. Normalization was done for SOZ and non-SOZ electrode populations separately within each time window. The ratio of SOZ over non-SOZ power in each respective decile at every time window was then calculated for input into a surgical outcome SRFE model. These ratios were first used to train six random forest models, consisting of 100 decision trees, with one random forest for each frequency band. Following this, an SRFE model (SRFE-*outcome*) was constructed incorporating the six-frequency range random forest models. Again, SRFE hyperparameters included a GLM as the metalearner algorithm, now with the lambda parameter set to 0.1 and the alpha value set to 0.5 (specifying a combination of LASSO and Ridge regression), to prevent over-regularization of the already normalized features.^33, 34^ The SRFE-*outcome* model was tuned by selecting a threshold to maximize model accuracy. For all cross-validation procedures, a leave-one-out patient cross-validation method was used, along with majority voting (≥ 50%) across multiple seizures for each patient to finalize predictions.

### FREEZ module: a toolbox for visualization of correlation analysis between spectral features and the SOZ

The different processes described in the previous subsections have been integrated into the FREEZ module GUI (<u>F</u>requency <u>R</u>ange <u>E</u>xplorer <u>E</u>pileptogenic <u>Z</u>one) which is freely available for visualization and to support the reproducibility of this study’s results. FREEZ is coded in R and R Shiny, enabling graphical user interface via any web browser, and created using the **R**eproducible **A**nalysis and **V**isualization of intracranial **E**EG (RAVE) platform.^28^ RAVE integration allows users to run on laptops, desktops, or in the cloud, with seamless integration between platforms as all module interaction is performed through a web browser. Because all computing in RAVE can be performed on any server or user machine, interaction with the module does not require independent computation.^35^ iEEG data set have to be preprocessed in RAVE 2.0 as described previously for further processing in FREEZ. **Fig. 2** shows the workflow of the FREEZ module. The open-source nature of FREEZ democratizes access to advanced epilepsy analysis tools, bridging gaps in epilepsy surgery referrals and promoting informed clinical decisions.^36–38^ Through both a web app and a Docker image, the tool allows users to interact with study results without requiring local software or patient data downloads, fostering reproducibility and wide accessibility. Finally, FREEZ allows to pipeline the whole workflow through a script in R. Please refer to ***Supplementary Methods*** for detailed module description and pipelining code example.

**Figure 2:**
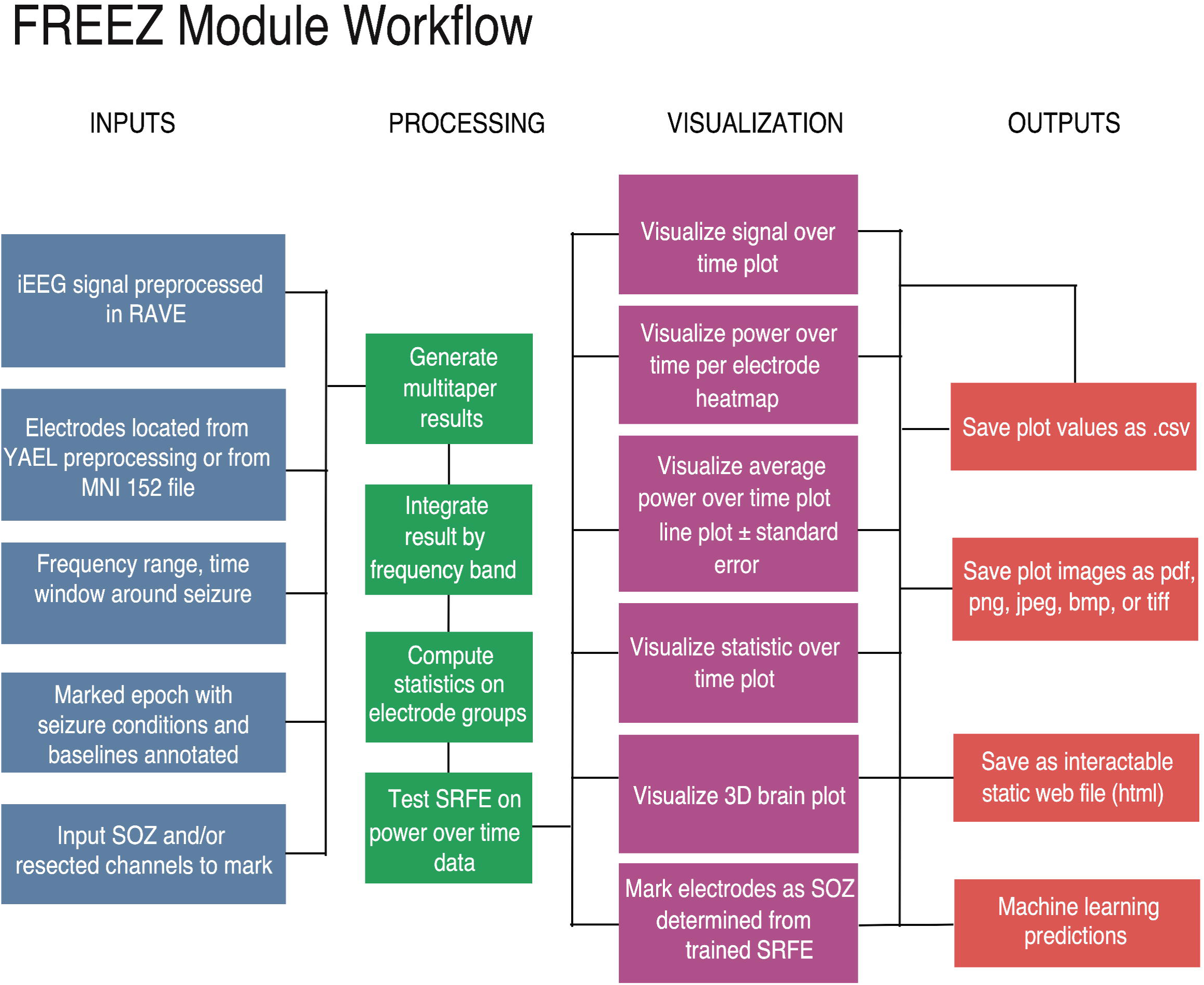
FREEZ module workflow. Data and user defined inputs shown in blue, intermediary processes in green, module visualization options in pink, and outputs in orange.

### Code and data Availability

Users can explore the FREEZ module GUI and the results for patient PT01 through a web application. They can also download a FREEZ module docker ^20^ with which can be run on PT01 or on their own patient data after preprocessing in RAVE. Please visit https://www.utmb.edu/neurosurgery/research/karas-laboratory/current-projects/rave-cezia/freez for more details on FREEZ module tutorials, the web application address, updates, and source code. Users can reach out for help about installation and software use in a dedicated Slack support channel.

## Results

21 patients were included in the study. Mean age at epilepsy onset was 19.5 ± 12 years (range: 13-59), and the mean age at the time of surgery was 36.39 ± 12.13 years (range: 13-59). The mean epilepsy duration before surgery was 16.89 years (range: 4-41). The cohort included 9 males, 9 females, and 3 patients with unspecified sex. All patients underwent electrocorticography (ECoG) intracranial monitoring with subdural grid and strip electrodes, with an average of 3.05 ± 0.84 ictal events recorded per patient (range: 1-4). Surgical targets were predominantly in the temporal lobe (N = 17), followed by the frontal lobe (N = 2) and the parietal lobe (N = 2). Post-surgery, 14 patients achieved seizure freedom (Engel I), while 7 did not. Among the non-seizure-free patients, 3 were classified as Engel II, 1 as Engel III, and 3 as Engel IV. Seizure onset patterns were observed as follows: A alone (PT06, PT07, PT11, PT12, PT13, PT15, PT16, PT17, UMMC009, UMF001), A and H (PT03), A and E (PT10), B and C (PT01), B and E (PT02), A and D (JH101), A and B (PT14, JH105), and H alone (UMMC005). Full details can be seen in ***Table 1***.

**Table 1:**
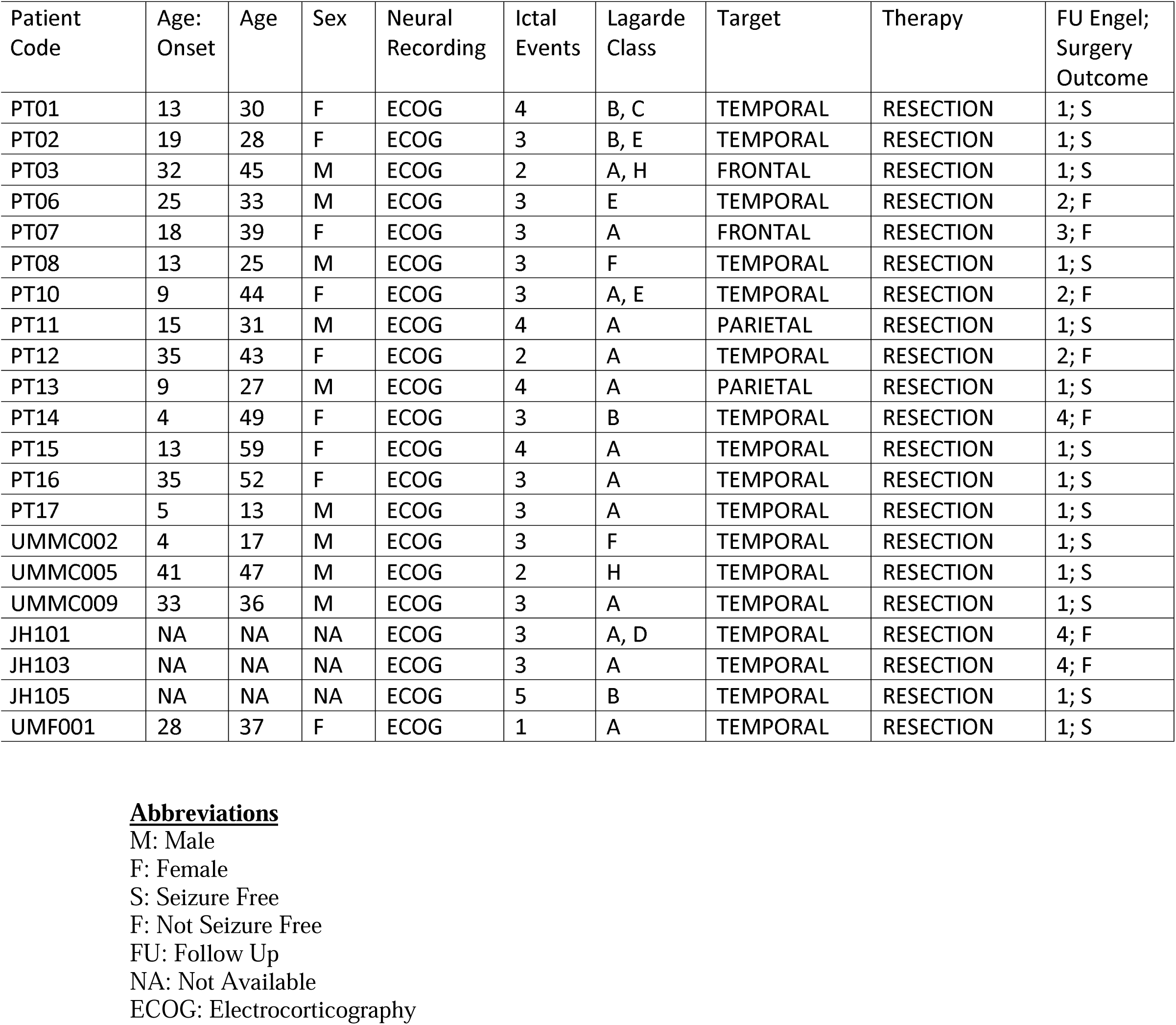
Characteristics of the included patients. Details include demographics, neural recording, surgical procedure, and outcome

| Patient Code | Age: Onset | Age | Sex | Neural Recording | Ictal Events | Lagarde Class | Target | Therapy | FU Engel; Surgery Outcome |
| --- | --- | --- | --- | --- | --- | --- | --- | --- | --- |
| PT01 | 13 | 30 | F | ECOG | 4 | B, C | TEMPORAL | RESECTION | 1; S |
| PT02 | 19 | 28 | F | ECOG | 3 | B, E | TEMPORAL | RESECTION | 1; S |
| PT03 | 32 | 45 | M | ECOG | 2 | A, H | FRONTAL | RESECTION | 1; S |
| PT06 | 25 | 33 | M | ECOG | 3 | E | TEMPORAL | RESECTION | 2; F |
| PT07 | 18 | 39 | F | ECOG | 3 | A | FRONTAL | RESECTION | 3; F |
| PT08 | 13 | 25 | M | ECOG | 3 | F | TEMPORAL | RESECTION | 1; S |
| PT10 | 9 | 44 | F | ECOG | 3 | A, E | TEMPORAL | RESECTION | 2; F |
| PT11 | 15 | 31 | M | ECOG | 4 | A | PARIETAL | RESECTION | 1; S |
| PT12 | 35 | 43 | F | ECOG | 2 | A | TEMPORAL | RESECTION | 2; F |
| PT13 | 9 | 27 | M | ECOG | 4 | A | PARIETAL | RESECTION | 1; S |
| PT14 | 4 | 49 | F | ECOG | 3 | B | TEMPORAL | RESECTION | 4; F |
| PT15 | 13 | 59 | F | ECOG | 4 | A | TEMPORAL | RESECTION | 1; S |
| PT16 | 35 | 52 | F | ECOG | 3 | A | TEMPORAL | RESECTION | 1; S |
| PT17 | 5 | 13 | M | ECOG | 3 | A | TEMPORAL | RESECTION | 1; S |
| UMMC002 | 4 | 17 | M | ECOG | 3 | F | TEMPORAL | RESECTION | 1; S |
| UMMC005 | 41 | 47 | M | ECOG | 2 | H | TEMPORAL | RESECTION | 1; S |
| UMMC009 | 33 | 36 | M | ECOG | 3 | A | TEMPORAL | RESECTION | 1; S |
| JH101 | NA | NA | NA | ECOG | 3 | A, D | TEMPORAL | RESECTION | 4; F |
| JH103 | NA | NA | NA | ECOG | 3 | A | TEMPORAL | RESECTION | 4; F |
| JH105 | NA | NA | NA | ECOG | 5 | B | TEMPORAL | RESECTION | 1; S |
| UMF001 | 28 | 37 | F | ECOG | 1 | A | TEMPORAL | RESECTION | 1; S |

### High power in alpha, beta, gamma and high-gamma bands correlate with the SOZ

We first inspected individual patients heatmaps, sorting them by SOZ vs non-SOZ electrodes, and observed higher power among SOZ electrodes (***Fig. 1B***). To determine whether this trend held across all patients, we calculated the AUC for mean power across patients for each frequency band **(*Fig. 1E*).**

In the patient population that became seizure free after surgery, we compared the AUC of the mean power over time in the first 20 seconds after seizure onset for SOZ *vs.* non-SOZ electrodes. AUC of mean power for SOZ electrodes was significantly higher compared to non-SOZ electrodes in alpha, beta, gamma, and high-gamma bands (***Fig. 3***. alpha 3.3 ± 5.8 e+10 SOZ *vs.* 0.2 ± 0.4 e+10 non-SOZ, p = 0.0272; beta 1.4 e+10 ± 2.4 e+10 *vs.* 0.05 ± 0.10 e+10, p = 0.0263; gamma 6.9 ± 9.2 e+08*vs.* 3.6 ± 4.9 e+07, p = 0.0013; and high gamma 2.4 ± 3.2 e+07 *vs.* 3.6 ± 10 e+06, p = 0.0086), but not for delta and theta bands (delta 9.3 ± 22.0 e+10 *vs.* 4.9 ± 11.0 e+10, p = 1.0; theta 6.0 ± 11.0 e+10 *vs.* 0.6 ± 1.0 e+10, p = 0.0955) (***Sup. Table 1***). This result suggests that increases in alpha, beta, gamma, and high-gamma mean power immediately after seizure onset correlates with the SOZ.

**Figure 3:**
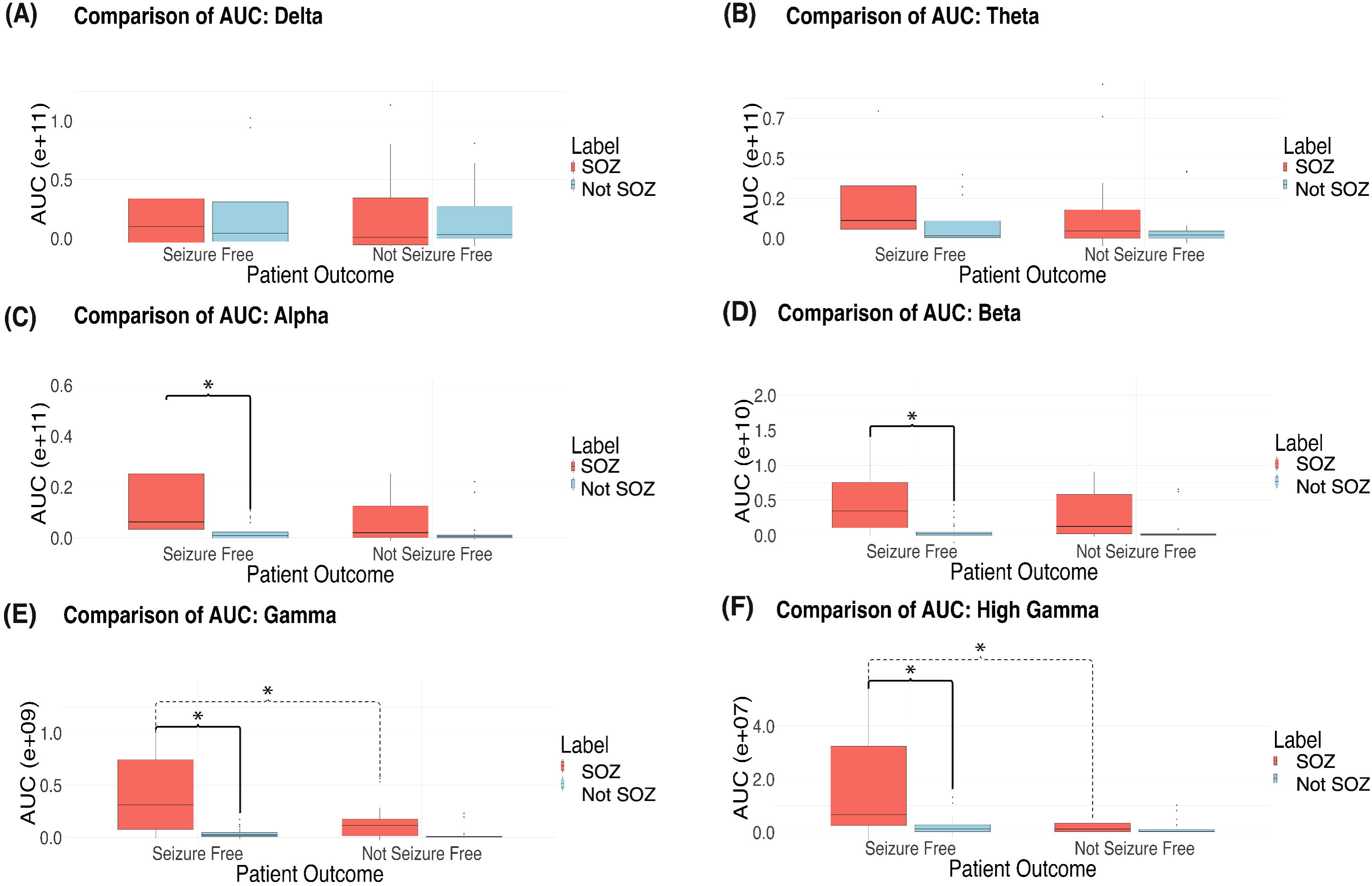
Area under the curve (AUC) for mean power over time from 0 to 20s following seizure onset. Seizure free and non-seizure free patients separated by marked SOZ and non SOZ channels in the delta (3A), theta (3B), alpha (3C), beta (3D), gamma (3E), and high gamma (3F) ranges. Statistical test used were two-sample t-test with Bonferroni correction (24 comparisons). Box-plots represent group-level averages of seizure AUCs (n = 14 seizure-free, n = 7 not seizure-free). Significant (P < 0.05 following Bonferroni correction 24 comparisons) pairwise comparisons marked with *.

We then performed the same analysis in the patient population that did not become seizure-free after surgery. While there were similar trends, the AUC of mean power for SOZ vs. non-SOZ electrodes did not reach statistical significance after correcting for multiple comparisons (***Sup. Table 1***).

Additionally, we compared AUC of mean power for SOZ electrodes in patients who were seizure-free *vs.* not seizure-free after surgery (***Fig. 3***). SOZ electrodes in seizure free patients had significantly higher AUC of mean power compared to in non-seizure free patients in the gamma (6.9 ± 9.2 e+08 seizure free *vs.* 1.4 ± 1.7 e+08 non-seizure free, p = 0.0145) and high gamma (2.4 ± 3.2 e+07 *vs.* 2.1 ± 2.4 e+06, p = 0.0024) bands, but not delta, theta, alpha, or beta bands (delta 9.3 ± 22.0 e+10*vs.* 0.56 ± 4.2 e+10, p = 0.8678; theta 6.0 ± 11.0 e+10 *vs.* 1.6 ± 2.7 e+10, p = 0.4956; alpha 3.3 ± 5.8 e+10 *vs.* 0.64 ± 0.86 e+10, p = 0.1320; and beta 1.4 ± 2.4 e+10 *vs.* 0.29 ± 0.31 e+10, p =0.1624) (***Supp. Table 2***). These results may highlight that the magnitude of the power difference is important and could reflect differences in SOZ sampling or underlying physiology that may affect seizure-freedom outcomes.

Finally, we compared AUC of mean power for non-SOZ electrodes in patients who became seizure-free *vs*. not seizure-free after surgery. There were no significant differences between these groups (***Sup. Table 2***). We then calculated the average power over frequency, instead of time, to better visualize and confirm differences between seizure-free and non-seizure-free SOZ contacts, as shown in ***Fig. 4***.

**Figure 4:**
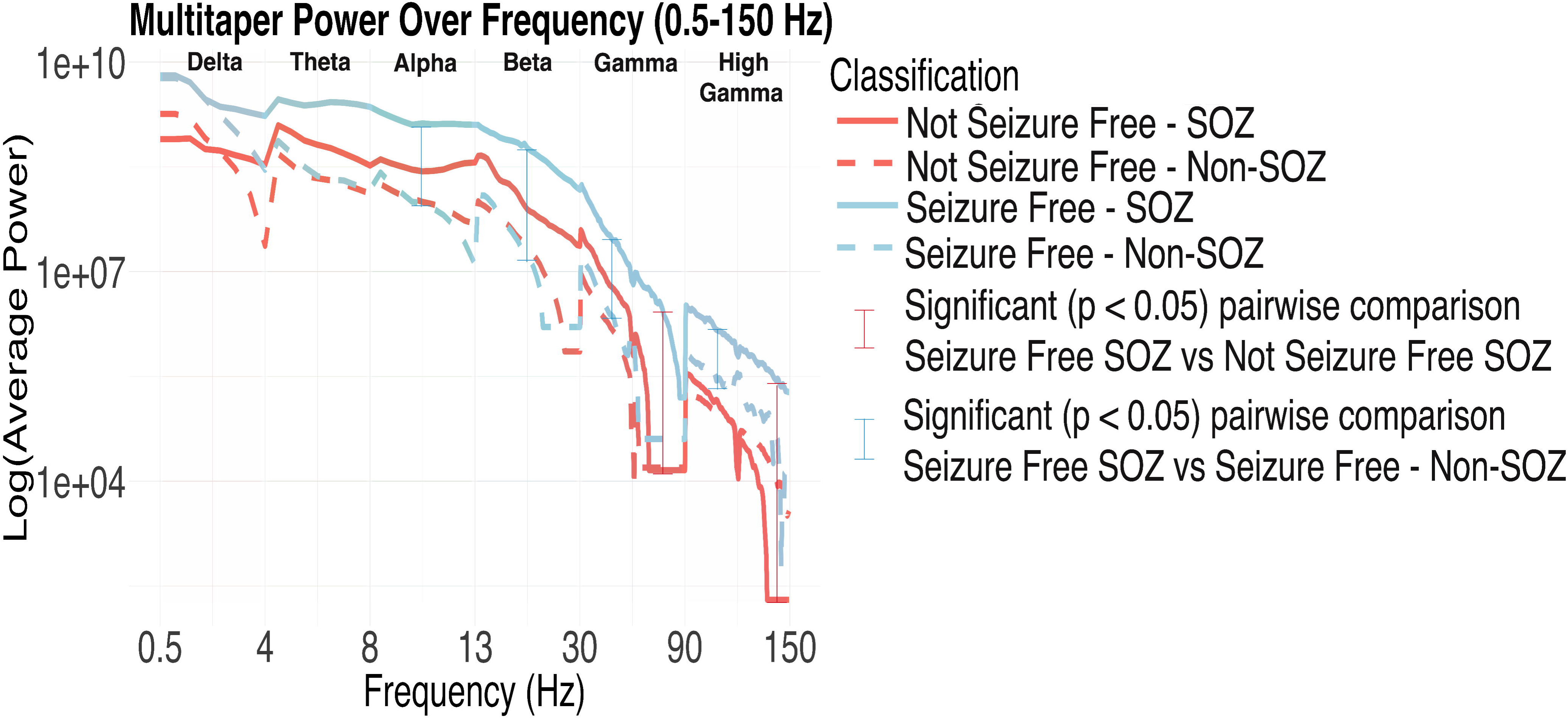
Multitaper power spectrum from 0.5 to 150 Hz, averaged over the first 20 seconds post-seizure onset, for four groups: seizure-free SOZ (solid blue), seizure-free non-SOZ (dashed blue), not seizure-free SOZ (solid red), and not seizure-free non-SOZ (dashed red). The Y-axis shows the scaled log10 of the average power, while the X-axis represents continuous frequencies evenly spaced between six defined bands: delta, theta, alpha, beta, gamma, and high gamma. Statistical comparisons between groups were performed using two-sample t-tests on AUCs for each band with Bonferroni correction (24 comparisons). Red brackets highlight significant pairwise comparisons (p < 0.05) between seizure-free SOZ and not seizure-free SOZ, and the blue bracket marks a significant comparison (p < 0.05 from two-sample t-tests on AUCs for each band) between seizure-free SOZ and seizure-free non-SOZ. Data shown are means ± SEM.

To confirm that observed differences in spectral power were not attributable to disparities in the number of SOZ-labeled electrodes across outcome groups, we additionally performed a two-sample t-test comparing the distribution of SOZ channel counts between seizure-free and non-seizure-free patients. Seizure-free patients (n = 14) had a mean of 14.5 ± 10.8 SOZ electrodes, while non-seizure-free patients (n = 7) had a mean of 11.0 ± 4.7 electrodes. This difference was not statistically significant (p = 0.316).

### Combining Spectral Power Bands with Machine Learning Improves Electrode Classification

With the above results suggesting that multiple different spectral power bands correlate strongly with the SOZ, we hypothesized that combining results across all power bands would improve SOZ localization. We therefore trained an SRFE machine learning model (SRFE-*electrode*) on the post-operative seizure-free cohort to integrate features across all power bands and identify electrodes within the SOZ. We justified the use of an SRFE model by determining that the predictions of individual base models across power bands are not highly correlated, suggesting an ensemble model would combine features effectively (***Sup. Fig. 1A***). Additionally, SRFE-*electrode* outperformed both the simple random forest and averaged random forest models (***Sup. Fig. 2***). An 86.07% threshold on the prediction probability output of SRFE-*electrode* was identified from leave-one-out cross validation to result in the greatest number of true positives while maintaining PPV ≥ 95% (***Fig. 5A***). In this cohort there were a total of 951 electrodes across 14 patients. 202 (21.2%) were marked SOZ by an epileptologist. SRFE-*electrode* yielded 28 true positive predictions within the SOZ and one false positive (***Fig. 5B***). These true positive electrodes were identified in 9 of the 14 (64%) seizure-free patients (PT01, PT02, PT08, PT13, PT15, PT17, UMMC002, UMMC005, JH105); the model did not identify any SOZ electrodes in 5 patients. SRFE-*electrode* achieved an accuracy of 81.6%, with a PPV of 96.6%, NPV of 81.1%, sensitivity of 13.9%, and specificity of 99.9%.

**Figure 5:**
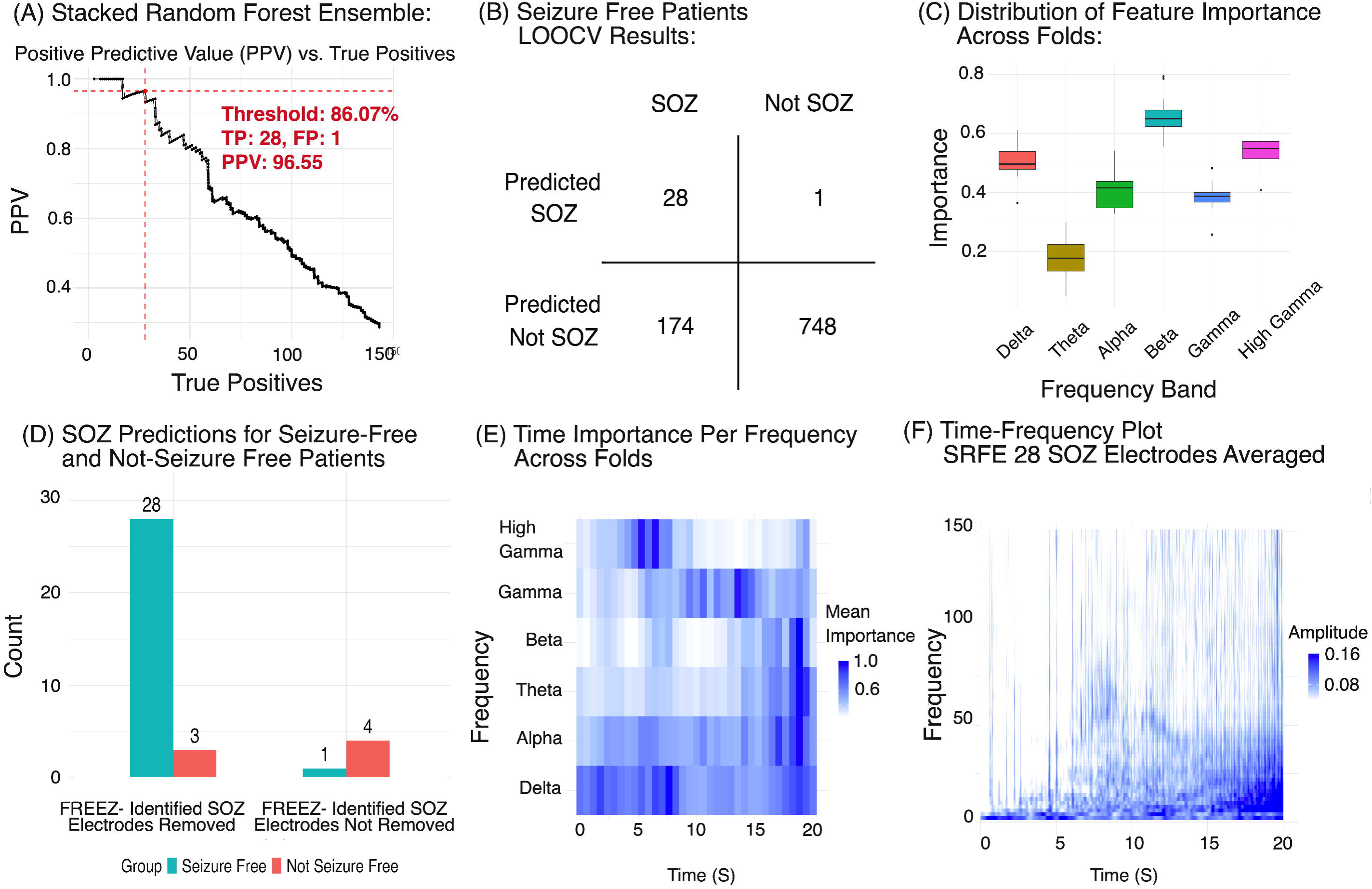
Machine learning results from the stacked random forest ensemble (SRFE). (5A) Positive predictive value (PPV) vs. true positives (TP) for changing threshold values (shown in blue text), with maximum true positives with PPV ≥ 95% marked by horizontal and vertical dotted red lines, annotated with threshold, TP, false positives (FP), and PPV. (5B) Results of leave-one-out patient cross-validation (LOOCV). (5C) Distribution of feature importance for the SRFE showing meta-learner coefficient for each base frequency band model. (5D) Results of SOZ predictions applied to non-seizure free patients showing distribution between resected and non-resected tissue, juxtaposed against seizure-free patients. (5E) Distribution of feature importance over time for random forest models trained on each frequency range. (5F) Average time-frequency plot from seizure onset to 20s following seizure onset for the 28 electrodes identified as SOZ by the SRFE model.

To gain insight on the contribution of each power band to SRFE-*electrode*’s SOZ classification, we extracted the GLM meta-learner’s optimized frequency-specific model coefficients. A larger base model coefficient indicates greater impact of that power band on the final SRFE-*electrode* model, identifying which of the six frequency bands most strongly influence SOZ determination. The beta power importance coefficient was highest (0.66 ± 0.07), followed by high gamma (0.54 ± 0.06), delta (0.51 ± 0.06), alpha (0.41 ± 0.07), gamma (0.38 ± 0.05), and theta range (0.18 ± 0.07) (***Fig. 5C***).

SRFE-*electrode* was then applied to the patient population that did not become seizure free after surgery (n=7). We hypothesized that the model would identify SOZ electrodes that had not been originally identified or removed. The model identified 3 out of 75 electrodes that had been labeled by clinicians as SOZ and removed, and 4 electrodes that were not labeled as SOZ and not removed (***Fig. 5D***). These electrodes were identified across 3 of the 7 non-seizure free patients.

Finally, we examined the temporal evolution of SRFE-*electrode*’s optimal weights by analyzing the variable importance derived from frequency-specific random forest models. Each model applied variable importance metrics to distinct frequency ranges across the time windows, allowing us to visualize the significance of each frequency band over a 20-second period (***Fig. 5E***). From seizure onset (0s) to 7.5s, we observed an initial phase dominated by delta activity, transitioning to a period of high gamma predominance between 5 and 7.5s. This was followed by a second phase, characterized by sustained gamma predominance from 7.5s to 15s. The third and final phase, spanning 15s to 20s, was marked by increased prominence in the theta, alpha, and beta frequency ranges, along with a resurgence in delta activity. We then correlated these findings with the averaged time-frequency plot for the 28 SOZ electrodes identified by the SRFE model, revealing that periods of high power in the time-frequency plots were temporally aligned with the random forest variable importance metrics (***Fig. 5F***).

### Combining Spectral Power Bands also Predicts Post-Operative Seizure Outcome

The second SRFE (SRFE-*outcome*) was trained using electrodes power deciles across spectral bands (***Fig. 6A***) to predict patient outcomes. Similar to SRFE*-electrode*, the Pearson correlation coefficients between power band predictions revealed low correlation justifying the choice of ensemble model (***Sup.*** Fig 1B). SRFE-*outcome* was also compared to predictions constrained to each frequency band individually. SRFE-*outcome* performed better on patient outcome prediction compared to any individual band RF (***Fig. 6B***). A 74.2% threshold for the SRFE-*outcome* model was identified from leave-one-out cross validation to result in the greatest number of true positives while maintaining PPV ≥ 95%. For the population of 21 patients, SRFE-*outcome* yielded 14 true positives, 6 true negatives, 1 false positive (PT12), and 0 false negatives (***Fig. 6C***). Overall, SRFE-*outcome* achieved an accuracy of 95.2%, with a PPV of 93.3%, NPV of 100%, sensitivity of 100%, specificity of 85.7%, and AUC of 0.91.

**Figure 6:**
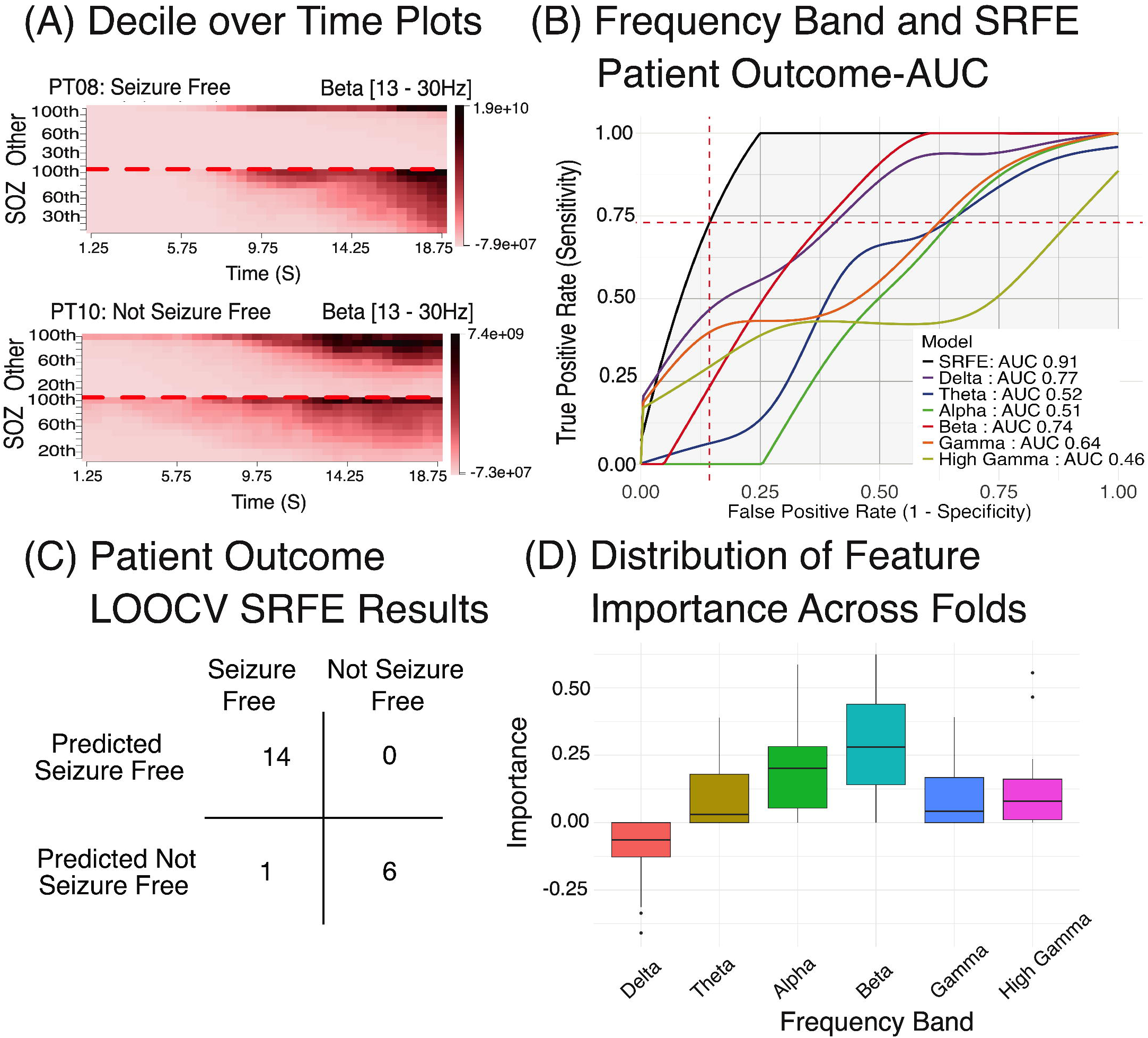
Results from patient outcome predictions are presented in four sections: 6A shows the statistics over time plots for a seizure free and non-seizure free patient in the beta frequency range [13 – 30 Hz]. 6B displays the Area Under the Curve (AUC) for individual frequency bands calculated using random forests, as well as the AUC for the final stacked random forest ensemble (SRFE). 6C presents the results from the leave-one-out patient cross validation at the maximum accuracy cutoff, which includes true positives, false positives, true negatives, and false negatives from the SRFE model. Finally, 6D shows the distribution of feature importance for the SRFE showing meta-learner coefficient for each base frequency band model.

To gain insight on the contribution of each power band to SRFE-*outcome*’s seizure outcome prediction, we extracted the model’s optimized frequency-specific random forest model coefficients. The beta power importance coefficient was highest (0.29 ± 0.07), followed by alpha (0.18 ± 0.15), high gamma (0.13 ± 0.15), theta (0.10 ± 0.13), and gamma (0.09 ± 11). Delta was determined to have a negative coefficient value (-0.10 ± 0.12) indicating an inverse correlation between the delta power decile feature and seizure outcome prediction utility. (***Fig. 6D***)

### FREEZ Software for Data Analysis and Visualization

FREEZ enables interactive analysis of data to examine results of SRFE-*electrode*. ***Fig. 7*** illustrates visualizations of SRFE-*electrode* results for four patients. It shows the heatmap results in the beta range. The heatmap displays the electrode numbers and the time in seconds, respectively on the y and x axes. Electrodes marked as SOZ are highlighted in blue and stacked on the heatmap bottom. Cross marks highlight electrodes that have been classified as SOZ by SRFE-*electrode* from the multi-patient analysis. ***Fig. 7A*** shows PT08, who was seizure-free, and had all predicted SOZ by SRFE-*electrode* resected. We can see on ***Fig. 7B*** another visualization of electrode classification result on PT15, a seizure free patient, with beta power outside of the resected SOZ. As such, SRFE-*electrode* selected electrodes both within and outside of resected tissue, resulting in the one false positive result. PT10 demonstrates the predictive power of SRFE-*electrode* on a non-seizure free patient, with the expected SOZ determined by the model to be outside of the tissue that was resected. This can be seen by the SOZ electrodes on the heatmap bottom that do not correlate well with high relative power in the area predicted by SRFE-*electrode* in ***Fig. 7C***. Finally, PT06 represents a patient where potentially suboptimal electrode implantation resulted in relative spectral power not being predictive of which electrodes should be removed. High relative power is within the removed electrodes and correlates with SRFE-*electrode*’s predictions as seen in ***Fig. 7D***, however the patient was not seizure free postoperatively.

**Figure 7:**
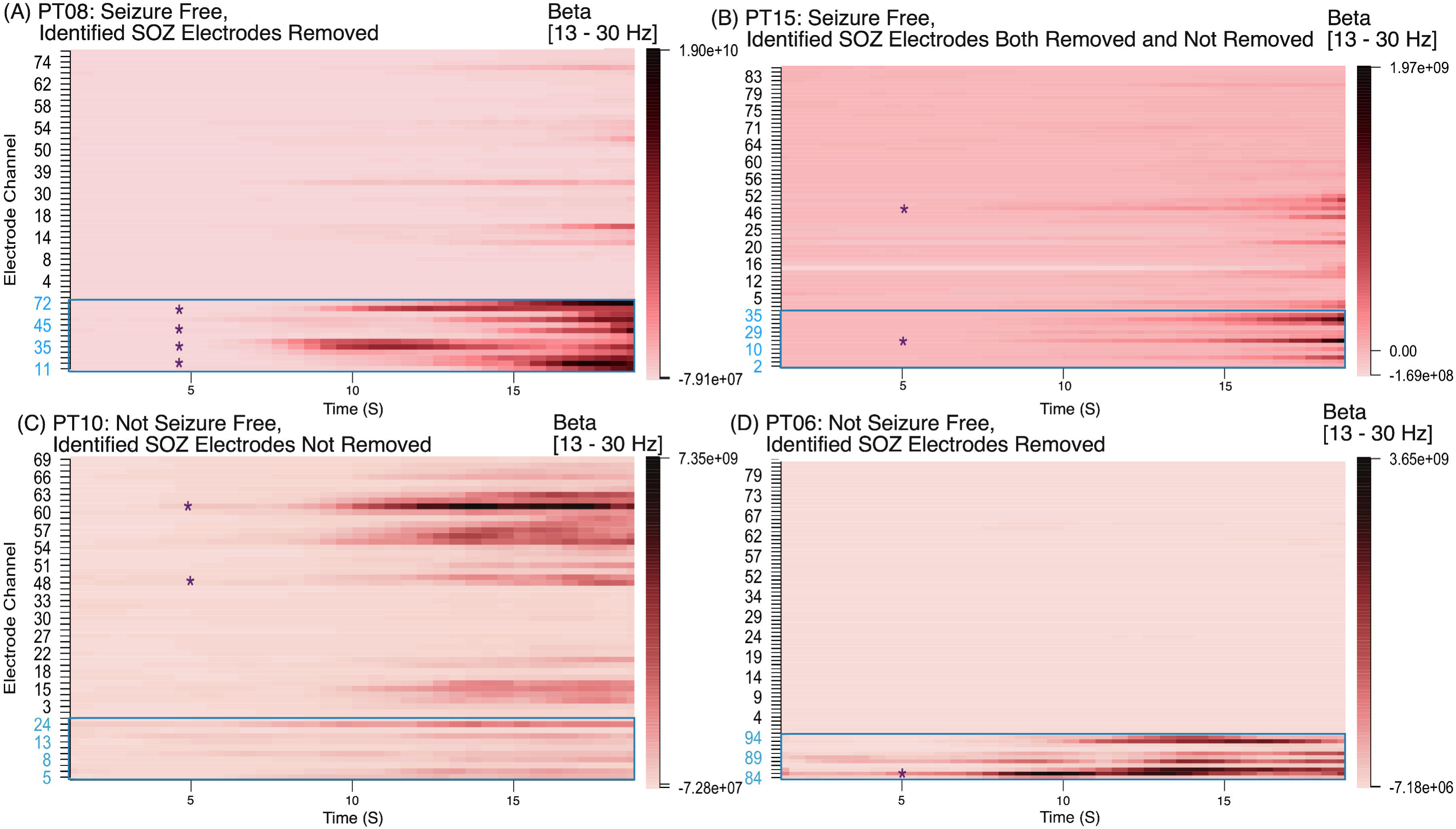
Patient multitaper heatmap results in the beta (13 – 30 Hz) range. SOZ channels (marked blue) are organized at the bottom of the heatmap and delineated with a blue box. Predicted SOZ channels by the random forest are highlighted with an asterisk (*). (7A) PT08, who was seizure-free, had all predicted SOZ resected. (7B) PT15, also seizure-free, had one predicted SOZ channel removed, and one not removed. (7C) PT10, who was not seizure-free, had predicted SOZ not removed. (7D) PT06, also not seizure-free, had the predicted SOZ removed.

The FREEZ software enables spatiotemporal visualization of the mean relative power across electrodes overlaid on patient brain models, implementation of SRFE-*electrode*, and script-based pipelining to generate results and provide statistical analysis and prediction using ensemble machine learning stacking models built on multi-frequency range analysis (***Sup. Fig. 3***).^19^

## Discussion

Accurate identification of the SOZ remains a challenge in the management of drug-resistant epilepsy. In this study, we characterize the power of multiple frequency bands at seizure onset in the SOZ and introduce an interpretable machine learning approach that integrates power spectra across a broad range of frequency bands within an SRFE model. This multi-band, ensemble-based strategy enhances the precision of both SOZ electrode classification and seizure outcome prediction, in contrast to only outcome prediction,^8, 10, 11, 14, 15^ providing a more comprehensive framework for interpreting the spectral dynamics of seizure onset regions. We emphasize the interpretability of this machine learning model for two reasons. First, we are reassured that the model labeling is utilizing meaningful characteristics of the iEEG signal. More strikingly, the weights of feature importance over time (***Fig. 5E***) are similar to the SOZ fingerprint^15^ reinforcing this fundamental neurophysiologic signature of the SOZ.

### Seizure Onset Zone Identification and Patient Outcome Prediction

We found a strong correlation between the SOZ and increases in alpha, beta, gamma, and high-gamma mean power at seizure onset in patients who became seizure free after surgery. While this finding is supported by prior literature,^8, 9, 15, 18, 28^ we also know that in a minority subset of patients early delta power is a marker of the SOZ. Notably DC shift is a known marker of the SOZ,^16, 39^ but are below 0.5 Hz and are therefore not fully reflected in the delta band. Additionally, other factors may contribute to scenarios when these findings are lacking in the SOZ. Less increase in mean power could be a result of poor sampling (e.g. incorrect pre-implantation hypothesis the SOZ location led to the SOZ being poorly sampled or not sampled). Alternatively, less increase in mean power could be due to a physiologic or genetic difference in the underlying tissue that is less amenable to surgical resection. To minimize these uncertainties, we utilized an interpretable machine learning approach to identify electrodes more accurately within the SOZ. The accuracy of these SRFE models suggest they are a relevant tool to provide a unified exploration of the correlation between spectral features and SOZ localization to help define the SOZ.

To reflect clinical workflows and enhance interpretability, we structured our pipeline into two distinct models: SRFE-*electrode*, which classifies electrodes as within or outside the SOZ, and SRFE-*outcome*, which predicts seizure freedom based on the spatial relationship between the proposed resection and SRFE-*electrode* labeled SOZ electrodes. This separation mirrors the clinical sequence of first identifying the SOZ and then considering its potential for resection and prognostic implications. Importantly, this modular design allows clinicians to independently interpret SOZ localization and outcome predictions, which is particularly useful in cases where the SOZ overlaps eloquent cortex or cannot be completely resected. By disentangling these steps, our approach enables more transparent decision-making and offers potential for integration with surgical planning (***Sup. Fig. 4***).

Our SFRE-*outcome* seizure predictions showed similar performance to other modern algorithms, for example the manifold random forest trained on the fragility method spatiotemporal results (AUC score 0.91 *vs.* 0.89).^8^ In addition to patient outcome prediction, automatic electrode classification is important to identify the SOZ. Our SFRE-*electrode* model performance compares favorably with prior work, for example the support vector machine trained on fingerprint features (PPV 96.6% *vs.* 90%).^40^ It is important to note, however, that the reported AUC and PPV values from the fragility and fingerprint methods were derived from separate datasets and are not directly comparable, instead providing general performance benchmarks.

The SRFE-*electrode* model was highly successful making true positive predictions for electrode classification even with heterogeneous seizure onset patterns. The percentage of patients with slow onset patterns, in the absence of fast activity, has been estimated at 20-25%. Previously described localization methods have difficulty identifying slow-onset patterns.^18, 41, 42^ 6 out of 21 patients (28.6%) included in this study fall into this slow-onset seizure pattern, including Lagarde et *al.* onset pattern E (PT02, PT06), F (PT08), E/F (PT10), and H (PT03, UMMC005). All slow-onset pattern patients were correctly predicted for seizure outcome. For electrode classification, all slow-onset electrodes were correctly labeled with two caveats. The SRFE-*electrode* model labeled SOZ electrodes in PT10 (not seizure free) that had not been labeled as SOZ in the dataset, suggesting the SOZ may have been originally misidentified. The SRFE-*electrode* model identified one electrode in PT06 within the SOZ but the patient was not seizure free. We suspect the iEEG implantation/sampling was suboptimal for this patient.

### Insights into the Neurobiology of Epilepsy

By wrapping multiple random forest models, each that detects time-dependent patterns in a given frequency band, within an ensemble model that assigns weights to each band, FREEZ provides a highly interpretable, data-driven approach to evaluate the temporal spectral pattern of the epileptogenic zone. The distribution of frequency band feature importance in the SRFE-*electrode* model (***Fig. 5C***) indicates that the beta, delta and high gamma frequency range are the most heavily weighted features across patients, which parallels prior literature.^8, 16, 43^ This insight could be used, for example, to improve accuracy of the epileptogenic index (high frequency: Eβ + Eγ / low frequency: Eθ + Eα). Instead of weighing all frequencies equally one could apply different coefficients accounting for differences in frequency band importance.^14^

Analysis of individual frequency band random forest models revealed three distinct periods of early ictal spectral patterns (***Fig. 5F***). Period 1 (0-7.5s) was characterized by a predominance of first delta and then high-gamma frequencies. In Period 2 (7.5-15s), gamma frequencies became dominant. Period 3 (15-20s) showed increased importance of theta, alpha, and beta power, with a reemergence of delta frequencies. This pattern mirrors previously observed time-frequency signatures of the SOZ, suggesting a consistent underlying physiological process during the ictal period.^14, 40^

The predominant seizure onset pattern, marked by fast activity, was observed in 71.4% of patients, consistent with previous reports of 75–80%.^18, 41, 42^ High gamma activity in the initial seizure onset reflects synchronized neuronal firing within the seizure core, with paroxysmal depolarizing shifts triggering burst firing.^44^ Delta activity is suppressed as gamma activity dominates as the seizure continues due to decreased inhibition from slow-inhibitory GABAergic neurons, allowing fast-spiking inhibitory interneurons, such as parvalbumin-positive cells, to generate gamma oscillations through rapid firing and tight coupling with excitatory neurons.^15, 45–49^ Previous studies including the epileptogenic index^14^, fingerprint method^15^, and HFO^11, 50^ have identified this activity and used it as a biomarker for SOZ identification. This high-frequency synchronization supports localized cortical processing, containing seizure activity and facilitating the transition to seizure termination.^15, 45–49^ In the final period, our model highlights a broad relevance across theta, alpha, and beta bands, along with reemergent delta frequencies, potentially linked to post-inhibitory rebound bursting in pyramidal neurons.^51^ Following intense inhibitory input, pyramidal neurons may exhibit rebound excitation due to hyperpolarization-activated cyclic nucleotide-gated (HCN) channels and low-threshold T-type calcium channels.^51^ This rebound activity spans multiple frequency bands: theta and alpha mark the re-establishment of rhythmic activity and cognitive processing, while beta reflects sensorimotor engagement and the return of normal cortical function.^51^ Reemerging delta frequencies may indicate large-scale network re-synchronization as the brain restores homeostasis, with thalamocortical circuits facilitating the recovery of normal rhythms across these bands.^51^

In patients with slow-onset patterns without fast activity, the SRFE model detected the SOZ by heavily weighting delta power at initial seizure onset. Slow-onset patterns are hypothesized to result from network-organized SOZs, where seizure propagation depends on extensive connectivity rather than focal initiation.^28^ This network organization facilitates delta frequency oscillations, which are able to engage widespread cortical and subcortical areas via thalamocortical pathways.^52^ During seizures, alterations in thalamocortical interactions can enhance delta activity due to disrupted inhibitory control and increased synaptic strength within local circuits.^52^ This interaction creates a permissive environment for seizure propagation, where increased coupling between the thalamus and cortex supports synchronized firing across broader regions of the SOZ.^52^ These slow onset patterns have been described more frequently in extratemporal or operculo-insular epilepsies, where thalamocortical and network dynamics are more pronounced compared to mesiotemporal epilepsy and as a result achieving seizure freedom is often more challenging.^28, 53^ Our SRFE model’s ability to distinguish these slow-onset patterns is crucial, because in cases where epileptogenic network disconnection is feasible, surgical resection is able to achieve seizure freedom.

### Limitations and Future Directions

The results of this study are limited by the relatively small sample size (n=21) and retrospective nature of the dataset. Future studies should validate these findings on larger prospective patient cohorts, utilizing prospective analysis. Because the fragility multi-center dataset did not include post-operative T1 MRI scans, the precise resection volumes were unavailable. Therefore, we relied on the authors claim that all electrodes labeled SOZ were resected, and used these electrode labels as the SOZ ground truth for training and validating both the electrode classification and seizure outcome prediction models. In our current data set, only 4 patients were extratemporal. Curated datasets should include a larger representation of these patients in order to analyze the impact of more diverse population in SRFE training and performance.^28, 53^ All patients in this study had ECoG electrodes implanted—the model still needs to be validated for the stereo EEG implantation technique.

## Acknowledgements

We thank Katrina Reichardt for assistance with data acquisition.

