## Supplemental Figure 1 for "Integrating Data Across Oscillatory Power Bands Predicts the Seizure Onset Zone in Focal Epilepsies"

(A) Electrode Prediction Correlation Matrix

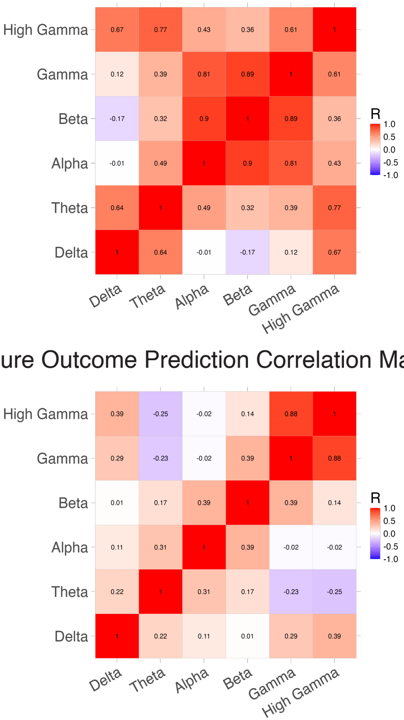


(B) Seizure Outcome Prediction Correlation Matrix



**Supplementary Figure 1:** Pearson correlation coefficients for machine learning outcome predictions. Correlations between power band predictions for electrode outcome predictions (A) and seizure outcome predictions (B).
