## Supplemental Figure 2 for "Integrating Data Across Oscillatory Power Bands Predicts the Seizure Onset Zone in Focal Epilepsies"

Stacked Random Forest Ensemble:

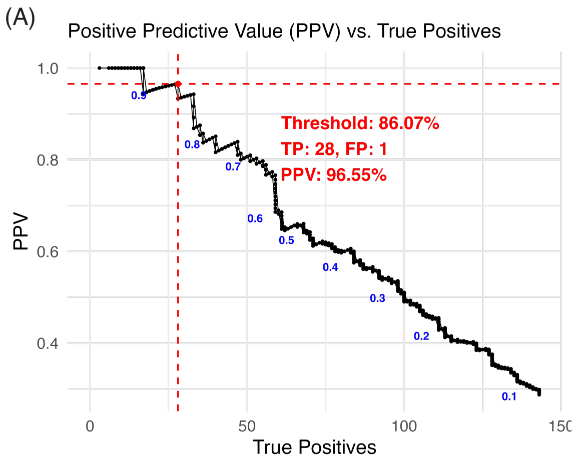


Random Forest:

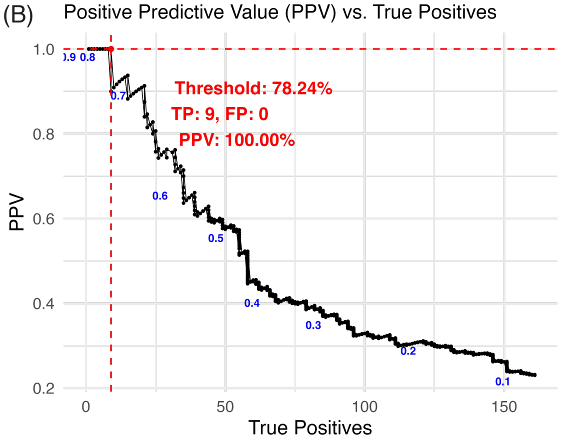


Averaged Random Forest:

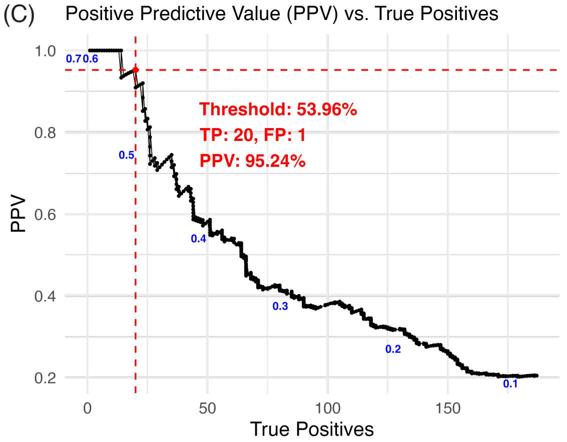


**Supplementary Figure 2:** Machine learning results on electrode selection for multiple models tested machine learning models. Positive predictive value (PPV) (y-axis) vs. true positives (x-axis) for three tested machine learning models on time-frequency features across six frequency ranges (delta, theta, alpha, beta, gamma, high gamma). (2A) Simple random forest model, (2B) frequency-specific random forests with stacked predictions, and (2C) stacked random forest ensemble (SRFE). Blue numbers represent applied thresholds to the output model probabilities. Dotted red lines mark the greatest number of true positives achieved with a PPV ≥ 95% considered the optimal threshold, with intersecting horizontal and vertical dotted red lines specifying the optimal threshold, as well as the number of true positives (TP), false positives (FP), and PPV at that optimal threshold.
