## Supplemental Figure 3 for "Integrating Data Across Oscillatory Power Bands Predicts the Seizure Onset Zone in Focal Epilepsies"

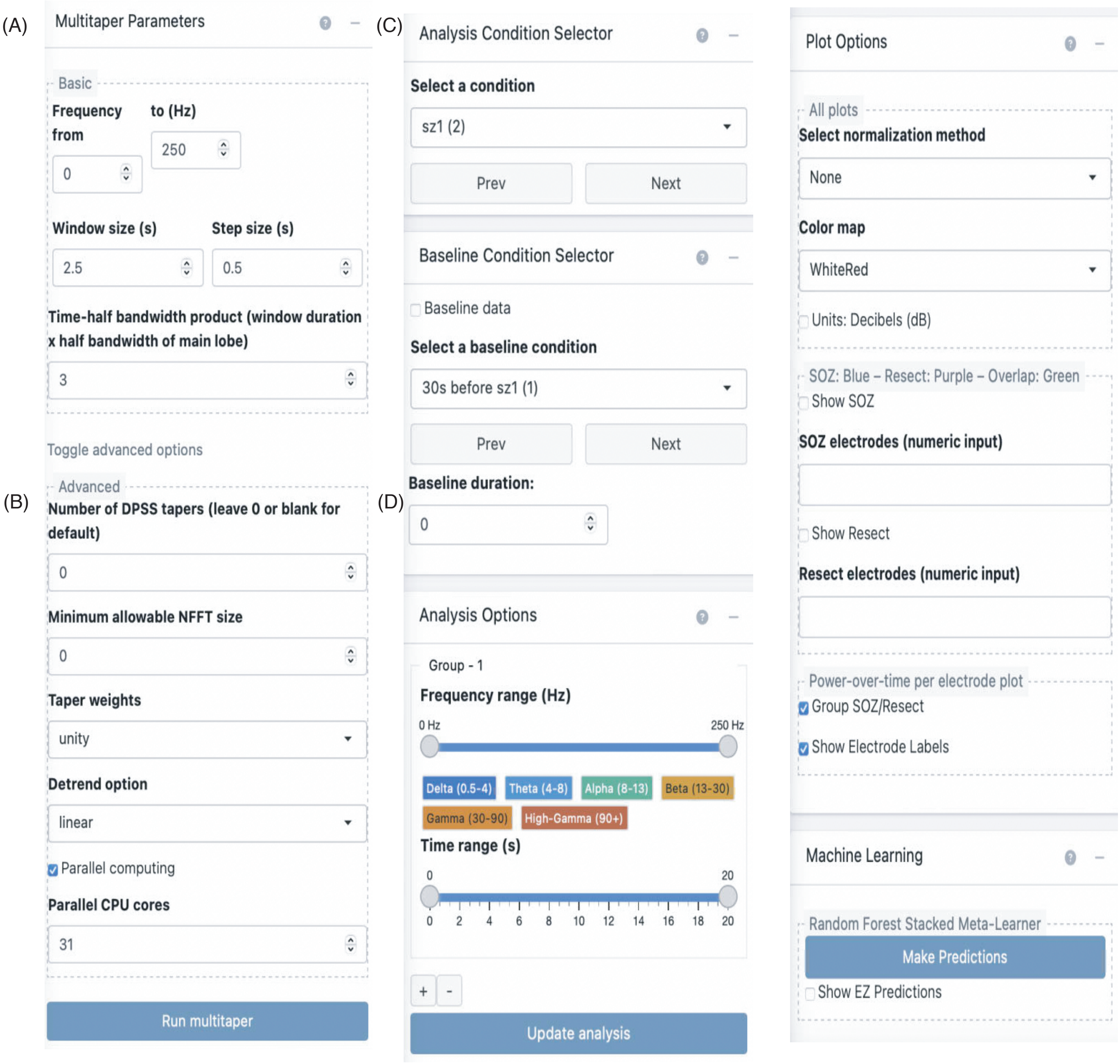


**Supplementary Figure 3:** Selection panels provided as user interface options in the FREEZ module. Basic parameter selection options include frequency range, window size, step size, and time-half bandwidth product (1A), while advanced options cover the number of DPSS tapers, minimum allowable NFFT size, taper weights, detrend options, and parallel computing options (1B). Analysis options allow for selection of baselining condition and duration (1C), and analysis frequency and time ranges, with default buttons for predefined ranges in delta (0.5-4 Hz), theta (4-8 Hz), alpha (8-13 Hz), beta (13-30 Hz), gamma (30-90 Hz), and high gamma (90+ Hz) (1D). Plotting options include selection of normalization method, color map, decibel units toggle, definition of seizure onset zone and resect electrodes (colored blue and purple respectively, with overlap in green), grouping of SOZ/Resect channels at the bottom of the map, and toggle for electrode labels (1E). Machine learning options include prediction capabilities and visualization on heatmaps (1F).
