## Supplemental Figure 4 for "Integrating Data Across Oscillatory Power Bands Predicts the Seizure Onset Zone in Focal Epilepsies"

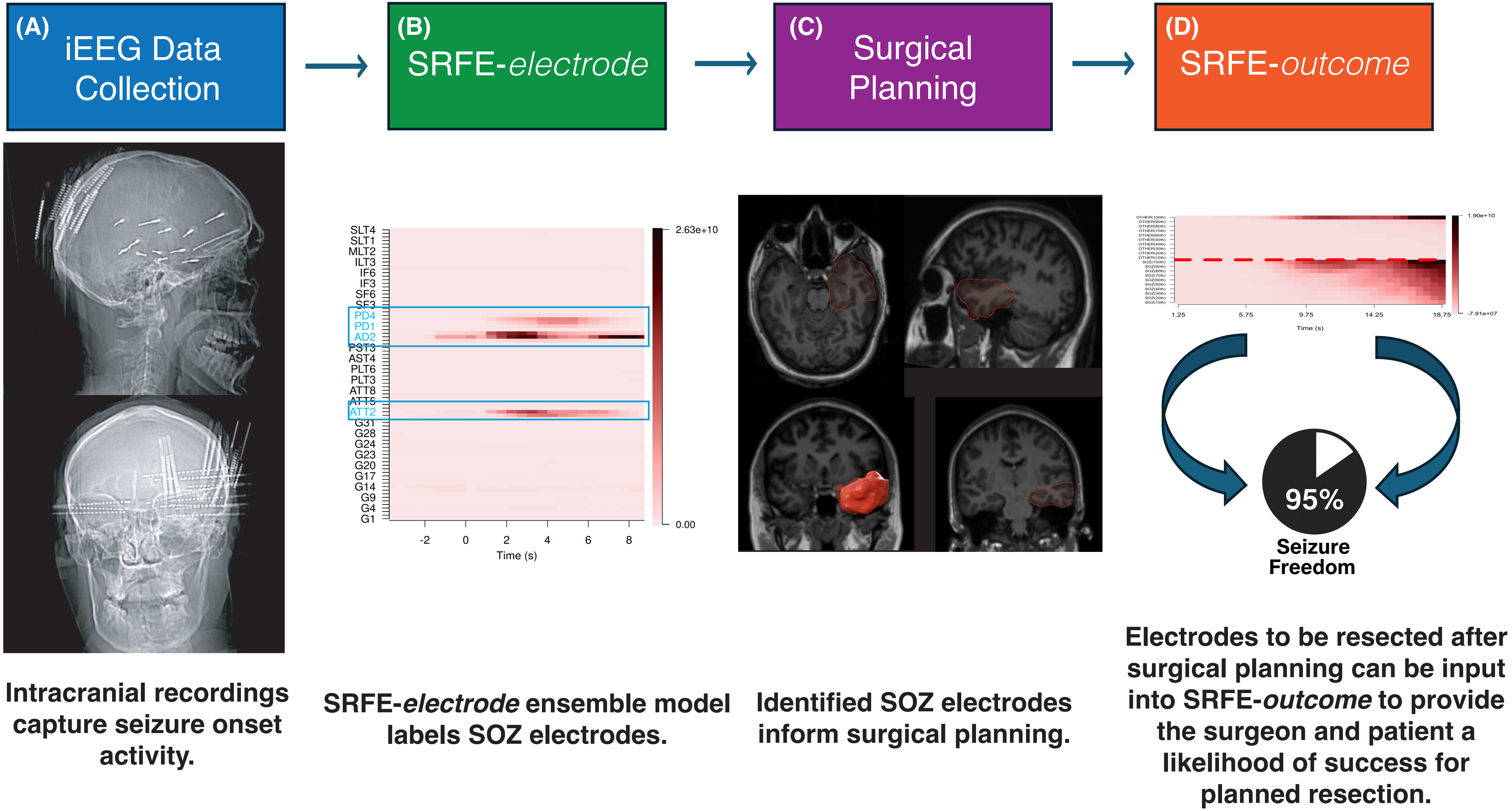


**Supplementary Figure 4:** End-to-end clinical workflow integrating FREEZ and SRFE models. Left-to-right: (A) iEEG data collection. Intracranial recordings capture seizure onset activity (example lateral and AP skull radiographs). (B) SRFE-electrode. Multitaper power heatmaps are analyzed by the stacked random-forest ensemble to label seizure-onset-zone (SOZ) electrodes; SOZ channels are outlined in blue. (C) Surgical planning. Identified SOZ electrodes inform proposed resection, visualized on patient MRI (example resection mask in red). (D) SRFE-outcome. The set of electrodes planned for resection is entered into the outcome model to estimate the probability of post-operative seizure freedom.
