## Supplemental Methods for "Integrating Data Across Oscillatory Power Bands Predicts the Seizure Onset Zone in Focal Epilepsies"

**SUPPLEMENTARY METHODS**

**Spectral power analysis with the multi-taper method**

To generate a time frequency plot with the multi-taper method, the basic parameters include the step size or window length for analysis with respective default values 0.5s and 2.5s. The main lobe's bandwidth, in turn, defines the frequency resolution or the smallest detectable frequency difference, and its default value is set to 3 for optimal resolution. Additionally, the time-half bandwidth product is determined by multiplying the window's length (N seconds) by half of the main lobe's bandwidth (BW/2). These values ensure accuracy for a wide range of frequencies^1,2^. Advanced customization choices include setting the number of DPSS tapers used for spectrogram computation, specifying the minimum allowable NFFT size for manual zero-padding adjustment to optimize the Fast Fourier Transform (FFT) computation, selecting taper weights (options include "unity," "eigen" for taper contribution weighted by eigenvalues, or "adaptive" to minimize broadband leakage), choosing a detrend method ("linear" for fitting a linear model and subtraction, "constant" for zero-mean data, or "off" for no detrending), and configuring the number of parallel CPU cores for processing.

**FREEZ module Graphic User Interface**

**Spatiotemporal analysis options**

The advanced parameters of a multi-taper spectral analysis (MSA) to compute mean power in a user-defined frequency range can be toggled in the FREEZ GUI (***Sup. Fig 3A and 3B)***. Following processing of the spatiotemporal power results, the user can choose the epoch time window around the seizure offset. The user can choose between several conditions of the RAVE preprocessed signal e.g. ictal signal of different recorded seizures, and toggle between conditions for analysis. The user can also mean subtract baseline the heatmaps to a selected baseline condition, with the option the duration from the start of the baseline condition that the user wishes to utilize, where *X* = *X* – mean (*X_B_*), where again *X* represents the original heatmap and *X_B_* the selected baseline condition for the selected baseline duration. The options for this are presented in ***Sup. Fig 3C***. Frequency ranges and time axis can be modified for the analysis condition, with the analysis options allowing for sliding bars to be utilized for tailoring the analysis results. We also provide buttons for preset frequency ranges of interest for user convenience encompassing Delta [0.5-4 Hz], Theta [4-8 Hz], Alpha [8-13 Hz], Beta [13-30 Hz], Gamma [30-90 Hz] and High Gamma [90+ Hz] frequency ranges. The user can also define additional analysis frequency ranges. The power over time plot as seen in ***Figure 1D*** can be seen jointly with the preprocessed signal result in ***Figure 1C*** for visualization of the spectral features extracted by the multitaper.

**Plotting options**

Plotting options are available [**see supplementary Figure 1E**]:

1/ The user can choose between a non-normalized spatiotemporal view, or a min-max normalization of the plot by time window. The user can also convert the power units (P) supplied by the multitaper result into decibels (dB), where dB=10 x log_10_(P). These options are all provided in ***Sup. Fig 1E***.

2/ The user can also input electrodes deemed SOZ or Resect for plotting. It resulted in the following color marking: SOZ coded blue, Resect Purple, and contacts with overlap between the input Resect and SOZ colored Green on the y axis of the spatiotemporal plots.

3/ Users can see either the electrode number from the channel files in the BIDS format or their name.

4/ Spatiotemporal visualization can be toggled to display electrodes grouped as resected/soz on the bottom and the remaining electrodes on the top. This can be useful to check visually that marked electrodes correlate with high relative power.

**Statistical results**

The module GUI provides also visualization of the statistical analysis of the mean power results by electrodes grouping using user input in the plotting options. FREEZ GUI offers the visualization of the average and standard error power over time plot within specified channel groups [***Figure 1E***]. Moreover, the user can visualize the quantile statistics over time plot, with electrode grouping separated with a red dotted line, which provides additional analysis of differences in the power between selected electrodes [***Figure 3F***].

FREEZ offers a similar brain visualization tool than YAEL(37). FREEZ provides the ability to visualize electrodes on a template, or patient specific brain model. Additionally, we provide an electrode projection functionality to take in account brain shape variability (see ***Figure 3A and 3B***). FREEZ allows patient specific visualization of brain cortex surface mesh in the FreeSurfer format^3^. This data can be computed using the FreeSurfer software and patient specific T1 MRI (magnetic resonance imaging) imaging. With FREEZ brain mapping functionality, the spectral power results can be projected over the patient specific or template 3D brain for visualization of anatomical position, as well as location of selected resect and soz channels, and temporal analysis through a sliding bar of changes in heatmap values over time allowing for viewing of the dynamic evolution of the heatmap, color coded based on the color template selected by the user [***Figure 3B***].

**Example Code for Pipelining**

To automate and streamline the analysis of iEEG data across multiple patients and frequency bands, we implemented a pipelined approach using the RAVE platform and the FREEZ module. The following code snippet provides an example of how the patient data is processed to extract relevant features, compute spectral power over different frequency bands, return machine learning results, and visualize the results through RAVE's plotting functions. This pipeline allows users to dynamically analyze multiple patients' iEEG data by automatically organizing results into frequency-specific folders and generating visual outputs for power analysis, ensuring a highly modular and reproducible workflow. Below, we outline key steps and describe their relevance to the FREEZ (Frequency Range Explorer Epileptogenic Zone) module.

Purpose and Structure of the Code

The code is designed to:

- Read patient data from a CSV file, which contains key variables such as patient IDs, seizure conditions, baselining condition, and electrode classifications (SOZ or non-SOZ).
- Loop through the data to extract power features from six frequency bands (delta, theta, alpha, beta, gamma, high-gamma) and apply spectral analysis.
- Automate visualization of spectral power features (e.g., power-over-time plots, line plots, and quantile plots) to assist in identifying the epileptogenic zone.
- Generate output folders dynamically for each patient and frequency band to save .CSV results for each plot in an organized manner, ensuring easy access to data for further statistical analysis.

Example data file and pipelining code can be found using the information and github link provided in our wiki page: <https://openwetware.org/wiki/Karas_Lab:FREEZ_Module>
