## Supplemental Tables 1-2 for "Integrating Data Across Oscillatory Power Bands Predicts the Seizure Onset Zone in Focal Epilepsies"

**Supplementary Table 1:** Area under the curve (AUC) average power for included patients in a time window from seizure onset to 20s following seizure onset. Data is mean baselined to a time window -30s to -20s before seizure onset. Reported P-Values for pairwise comparison between seizure condition SOZ AUC and seizure condition not SOZ AUC. P < 0.05 is significant following Bonferroni correction for multiple P-Value testing. * indicates significance.

| Seizure Free | | | |
| --- | --- | --- | --- |
| Frequency Range | Seizure Condition SOZ AUC (N = 41) | Seizure Condition Not SOZ AUC (N = 41) | |
|  | AUC Average Power ± SD (e+10) | AUC Average Power ± SD (e+10) | P-Value |
| Delta [0.5-4 Hz] | 9.3 ± 22.0 | 4.9 ± 11.0 | 1.0000 |
| Theta [4-8 Hz] | 6.0 ± 11.0 | 0.6 ± 1.0 | 0.0955 |
| Alpha [8-13 Hz] | 3.3 ± 5.8 | 0.17 ± 0.36 | 0.0272* |
| Beta [0-30 Hz] | 1.4 ± 2.4 | 0.053 ± 0.098 | 0.0263* |
| Gamma [30-90 Hz] | 6.9 ± 9.2 e-2 | 3.6 ± 4.9 e-3 | 0.0013* |
| High-Gamma [90-256 Hz] | 2.4 ± 3.2 e-3 | 3.6 ± 10.0 e-4 | 0.0086* |
| Not Seizure Free | | | |
| Frequency Range | Seizure Condition SOZ AUC (N = 20) | Seizure Condition Not SOZ AUC (N = 20) | |
|  | AUC Average Power ± SD (e+10) | AUC Average Power ± SD (e+10) | P-Value |
| Delta [0.5-4 Hz] | 0.56 ± 4.2 | 1.5 ± 2.5 | 1.0000 |
| Theta [4-8 Hz] | 1.6 ± 2.7 | 0.6 ± 1.3 | 1.0000 |
| Alpha [8-13 Hz] | 0.64 ± 0.86 | 0.25 ± 0.61 | 1.0000 |
| Beta [0-30 Hz] | 0.29 ± 0.31 | 0.077 ± 0.19 | 0.3710 |
| Gamma [30-90 Hz] | 1.4 ± 1.7 e-2 | 2.9 ± 6.4 e-3 | 0.2410 |
| High-Gamma [90-256 Hz] | 2.1 ± 2.4 e-4 | 1.6 ±2.9 e-4 | 1.0000 |

**Abbreviations**

SD: Standard Deviation

HZ: Hertz

**Supplementary Table 2:** Area under the curve (AUC) average power for included patients in a time window from seizure onset to 20s following seizure onset. Data is mean baselined to a time window -30s to -20s before seizure onset. Reported P-Values for pairwise comparison between seizure free AUC and not seizure free AUC. P < 0.05 is significant following Bonferroni correction for multiple P-Value testing. * indicates significance.

| Seizure Condition SOZ | | | |
| --- | --- | --- | --- |
| Frequency Range | Seizure Free AUC (N = 41) | Not Seizure Free AUC (N = 41) | |
|  | AUC Average Power ± SD (e+10) | AUC Average Power ± SD (e+10) | P-Value |
| Delta [0.5-4 Hz] | 9.3 ± 22.0 | 0.56 ± 4.2 | 0.8678 |
| Theta [4-8 Hz] | 6.0 ± 11.0 | 1.6 ± 2.7 | 0.4956 |
| Alpha [8-13 Hz] | 3.3 ± 5.8 | 0.64 ± 0.86 | 0.1320 |
| Beta [0-30 Hz] | 1.4 ± 2.4 | 0.29 ± 0.31 | 0.1624 |
| Gamma [30-90 Hz] | 6.9 ± 9.2 e-2 | 1.4 ± 1.7 e-2 | 0.0145* |
| High-Gamma [90-256 Hz] | 2.4 ± 3.2 e-3 | 2.1 ± 2.4 e-4 | 0.0024* |
| Seizure Condition Not SOZ | | | |
| Frequency Range | Seizure Free AUC (N = 20) | Not Seizure Free AUC (N = 20) | |
|  | AUC Average Power ± SD (e+10) | AUC Average Power ± SD (e+10) | P-Value |
| Delta [0.5-4 Hz] | 4.9 ± 11.0 | 1.5 ± 2.5 | 1.0000 |
| Theta [4-8 Hz] | 0.6 ± 1.0 | 0.6 ± 1.3 | 1.0000 |
| Alpha [8-13 Hz] | 0.17 ± 0.36 | 0.25 ± 0.61 | 1.0000 |
| Beta [0-30 Hz] | 0.053 ± 0.098 | 0.077 ± 0.19 | 1.0000 |
| Gamma [30-90 Hz] | 3.6 ± 4.9 e-3 | 2.9 ± 6.4 e-3 | 1.0000 |
| High-Gamma [90-256 Hz] | 3.6 ± 10.0 e-4 | 1.6 ±2.9 e-4 | 1.0000 |

**Abbreviations**

SD: Standard Deviation

HZ: Hertz
